# Completing missing MeSH code mappings in UMLS through alternative expert-curated sources

**DOI:** 10.1101/602540

**Authors:** Eduardo P. García del Valle, Gerardo Lagunes García, Lucia Prieto Santamaría, Massimiliano Zanin, Alejandro Rodríguez González, Ernestina Menasalvas Ruiz

## Abstract

The increasing availability of biological, clinical and literary sources enables the study of diseases from a more comprehensive approach. However, the interoperability of these sources, particularly of the codes used to identify diseases, poses a major challenge. Because of its role as a hub of multiple medical vocabularies, the Unified Medical Language System (UMLS) has become one of the most widely used resources for mapping diseases from different classification systems. The coverage of these mappings, nevertheless, is still limited, so researchers must resort to other methods to fill these gaps. In this article we analyze the limitations in UMLS mappings for the main disease vocabularies and propose the exploitation of some alternative expert-curated sources to complete it. As a result, we demonstrate that this approach allows resolving more than 50% of the missing mappings for some vocabularies. All the findings are shared for validation and reuse.

## I. INTRODUCTION

In 2016, the then House Majority Leader Kevin McCarthy claimed that “there’s 10,000 diseases, and we only have 500 cures”. The Washington Post Fact Checker decided to examine the accuracy of this cite and contrasted these numbers [1]. They found out that he might have come up with the “10,000 diseases” by going through the Orphanet data set, which then contained 9,235 orphan or “rare” diseases. Another plausible source could have been a World Health Organization (WHO) statement that “scientists currently estimate that over 10,000 of human diseases are known to be monogenic,” meaning involving a single gene. The same article compared the number with the 10th revision of the International Statistical Classification of Diseases (ICD-10) which contains nearly 70,000 codes.

This anecdotal research illustrates a widespread problem in the field of disease research: the existence of multiple vocabularies and codifications of diseases, and the difficulty in relating them. This variety is explained because each vocabulary was originally created to meet a specific need. ICD codes for instance are used by doctors, health insurance companies, and public health agencies across the world to represent diagnoses. Because ICD has been translated to many languages, it is useful for text mining of clinical narrative in EHRs. However, its structure and disease names are not suited for biomedical literature mining [2]. In contrast, the Medical Subject Headings (MeSH) vocabulary is especially employed for the purpose of indexing journal articles and books in the life sciences; it serves as a thesaurus that facilitates searching. It is used by the MEDLINE/PubMed article database, by the National Library of Medicine’s (NLM) catalog of book holdings, and also by ClinicalTrials.gov registry to classify which diseases are studied by trials registered in ClinicalTrials.

A newer alternative to ICD and MeSH is the Systematized Nomenclature of Medicine – Clinical Terms (SNOMED CT). It cross-maps to several revisions of ICD and has a considerably broader scope than just diseases. SNOMED CT is a clinical terminology designed to capture and represent patient data for clinical purposes. SNOMED CT enables information input into an EHR system during the course of patient care, while ICD facilitates information retrieval, or output, for secondary data purposes. SNOMED CT is owned by the International Health Terminology Standards Development Organization (IHTSDO), an international non-profit organization founded in 2007. Other widely used disease classification systems, although of more specific use, are OMIM (genetic disorders), Orphanet (rare diseases) or NCI (carcinogenic diseases).

For years, these classifications evolved independently, making their interrelation difficult. However, the growing number of disease studies based on the integration of different biological and literary sources entailed the need to establish a mapping between them. One of the most notable efforts in this direction is the Unified Medical Language System (UMLS). Created in 1986 and maintained by the NLM, UMLS is a compendium of many controlled vocabularies in the biomedical sciences. The Metathesaurus conforms the core of UMLS and as of version 2018AB it comprises over 3 million biomedical concepts. It provides a mapping structure among the different vocabularies and thus allows one to translate among the various terminology systems. As a result, numerous studies and tools have used UMLS as an authentic Rosetta stone of disease terms.

However, due to the disparity of scopes and granularity of the disease vocabularies, the annotation and mapping of UMLS terms is a complex and still unfinished task. In the last decades, several initiatives have tried to improve and complete the mapping of diseases between different vocabularies by diverse methods. Already in 1998, a study by Bodenreider et al. proposed to use the semantic relationships between concepts to map terms of different vocabularies in UMLS with MeSH terms [3]. A later investigation contemplated the use of drug prescriptions to complete the mapping between MeSH and ICD-10 terms extracted from the UMLS Metathesaurus [4]. In 2011, Brandt et al. presented a method for creating mappings between the Orphanet terminology of rare diseases and UMLS. This method was based in the aggressive normalization of terms and the semantic ranking of partial candidate mappings in order to group similar mappings and attribute higher ranking to the more informative ones [5].

More recently, the emergence of massive source integration projects has driven the search for solutions to unify concepts. To build Hetionet, a heterogeneous network with information on diseases extracted from 29 sources, Himmelstein et al. created a simplified version of the Disease Ontology (DO) composed of only 137 terms. The so-called *DO Slim* eliminated the hierarchy of terms and maintained only those specific enough to be clinically relevant and general enough to be well annotated. Then they used transitive closure to map external disease concepts onto DO Slim [6]. The Monarch Initiative, a collaborative effort that aims to semantically integrate genotype–phenotype data from many species and sources, created the Monarch Merged Disease Ontology (MonDO) to integrate multiple human disease resources into a single ontology using a Bayes ontology merging algorithm [7]. The MalaCards human disease database started from existing and widely used categorization systems like ICD-10 and Orphanet and applied an algorithm using a choice of category-specific keywords contained in disease names and annotations to connect diseases from different sources [8]. Only 2,282 entries are successfully unified among 3 or 4 of the sources [9].

Despite their improvement, the use of automated mapping techniques is subject to errors that can be propagated to subsequent studies [10, 11]. Therefore, expert validation is still imperative. As an alternative, this article proposes the exploitation of expert-curated sources to complete the mapping of diseases, avoiding error propagation and the need for additional validation. First, we analyze the mapping coverage of MeSH disease terms to ICD-10-CM and SNOMED CT in UMLS. As described in the Methods section, around 50% and 15% of the mappings are missing for these vocabularies, respectively. Next, we propose the use of the DO and the ICD-10 to SNOMED CT mapping project by the IHTSDO to complete the unavailable connections. Combining these sources, about 54% of the missing mappings for ICD-10-CM and 8% for SNOMED CT can be retrieved, as detailed in the Results section. Finally, we collect the data obtained in supplementary files as a contribution for future studies^1^.

## II. METHODS

### A. Analysis missing MeSH mappings to ICD-10 and SNOMED CT in UMLS

First, we analyzed the mapping coverage between MeSH disease terms and those of SNOMED CT and ICD-10 in UMLS. Since the last available version of UMLS is 2018AB, we consulted the MeSH resources corresponding to 2018 (ftp://nlmpubs.nlm.nih.gov/online/mesh/2018). In this version, MeSH contains 4,903 descriptors under the categories C (Diseases) and F03 (Mental Disorders). Some descriptors may belong to more than one category. Table I shows the number of descriptors for each first level subcategory.

**TABLE I.**
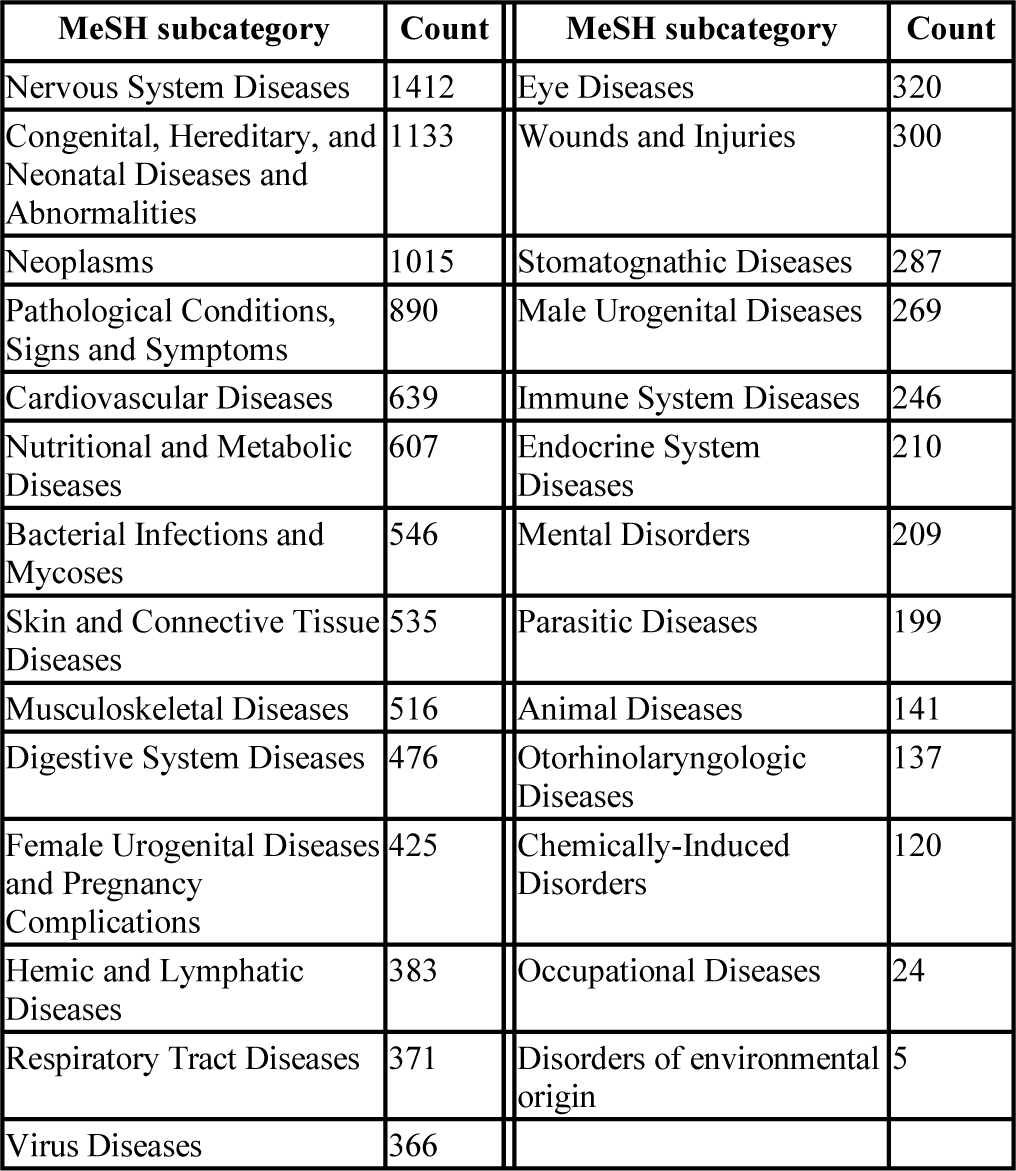
MESH DESCRIPTOR COUNT BY TOP LEVEL SUBCATEGORY (CXX AND F03)

| MeSH subcategory | Count | MeSH subcategory | Count |
| --- | --- | --- | --- |
| Nervous System Diseases | 1412 | Eye Diseases | 320 |
| Congenital, Hereditary, and Neonatal Diseases and Abnormalities | 1133 | Wounds and Injuries | 300 |
| Neoplasms | 1015 | Stomatognathic Diseases | 287 |
| Pathological Conditions, Signs and Symptoms | 890 | Male Urogenital Diseases | 269 |
| Cardiovascular Diseases | 639 | Immune System Diseases | 246 |
| Nutritional and Metabolic Diseases | 607 | Endocrine System Diseases | 210 |
| Bacterial Infections and Mycoses | 546 | Mental Disorders | 209 |
| Skin and Connective Tissue Diseases | 535 | Parasitic Diseases | 199 |
| Musculoskeletal Diseases | 516 | Animal Diseases | 141 |
| Digestive System Diseases | 476 | Otorhinolaryngologic Diseases | 137 |
| Female Urogenital Diseases and Pregnancy Complications | 425 | Chemically-Induced Disorders | 120 |
| Hemic and Lymphatic Diseases | 383 | Occupational Diseases | 24 |
| Respiratory Tract Diseases | 371 | Disorders of environmental origin | 5 |
| Virus Diseases | 366 |  |  |

Second, using the *Search* REST API of the UMLS Terminology Services (UTS), we checked that all 4,903 MeSH descriptors associated with diseases are contained in the UMLS, this is, they have a Concept Unique Identifier (CUI) in the Metathesaurus. Finally, we went through each MeSH descriptor (source path) and leveraged the UTS *Crosswalk* REST API to obtain their associated concepts in ICD-10-CM and SNOMED CT by setting the *targetSource* param to these vocabularies. We found that 2,444 MeSH descriptors correspond (i.e. have the same CUI) to 3,188 ICD-10-CM codes through 3,487 associations. Most missing mappings were found among *Neoplasms*, with 745 MeSH descriptors not mapped to any ICD-10-CM code for this top-level code. For SNOMED CT, we retrieved 4,199 MeSH descriptors associated to 7,769 SNOMED CT codes through 7,982 associations. In this case, most missing mappings affected to the *Pathological Conditions, Signs and Symptoms* type, with 163 MeSH descriptors not mapped to a SNOMED CT code.

### B. Completing missing UMLS mappings with the Disease Ontology

A frequently used alternative to UMLS to map disease vocabularies is the Disease Ontology (DO) [6,12–13]. This project is hosted at the Institute for Genome Sciences at the University of Maryland School of Medicine and was initially developed in 2003 to address the need for a purpose-built ontology that covers the full spectrum of disease concepts annotated within biomedical repositories in an ontological framework that is extensible to meet community needs [14]. Unlike UMLS, the DO project’s data is licensed under Creative Commons CC0, the most open license, to enhance collaboration and data sharing and to encourage broad and open usage. The latest DO release (GitHub, release 45, v2018-09-10) includes 9,069 DOID disease terms, with 62% of terms having a textual definition [15], and code mappings of 24 vocabularies, including SNOMED CT, UMLS, ICD-10-CM and MeSH. Table II contains the most represented vocabularies in the DO.

**TABLE II.** MOST REPRESENTED VOCABULARIES IN THE DISEASE ONTOLOGY

| Vocabulary | Total associated concepts | Unique vocabulary codes |
| --- | --- | --- |
| SNOMED CT | 12,443 | 11,476 |
| UMLS | 6,583 | 6,524 |
| NCI | 4,599 | 4,562 |
| ICD-10-CM | 4,226 | 2,909 |
| OMIM | 4,166 | 4,119 |
| MESH | 3,086 | 2,802 |
| ICD-9-CM | 2,626 | 2,591 |
| ORDO | 1,131 | 1,059 |
| EFO | 132 | 132 |
| KEGG | 39 | 37 |

First, we obtained the DO data from the project repository (http://disease-ontology.org/downloads) and extracted the terms that contained at least a MeSH descriptor code. We found 1,551 terms mapped to both MeSH and ICD-10-CM, and 2,457 terms mapped to both MeSH and SNOMED CT. Only 1,526 terms where mapped to MeSH, ICD-10-CM and SNOMED CT simultaneously. Then we compared the MeSH to ICD-10-CM and MeSH to SNOMED CT mappings in DO and UMLS, and obtained the symmetric difference of the two sets. In both cases, we found mappings in the DO that are not included in UMLS, as described in detail in the Results section.

### C. PubMed Completing missing UMLS mappings with the SNOMED CT to ICD-10-CM mapping project

A project started in 2011 by the WHO and the IHTSDO is building a SNOMED CT to ICD-10-CM map to support semi-automated generation of ICD-10-CM codes from clinical data encoded in SNOMED CT for reimbursement and statistical purposes [16]. Dual independent mapping by trained terminology specialists is employed to assure quality and reduce variability. Identical maps created independently are accepted as final, while discordant maps are reviewed by a third expert. Regular team meetings are held to discuss problematic and ambiguous cases.

The NLM distributes this map under the same license of use of the UMLS Metathesaurus. The map contains 123,260 unique SNOMED CT codes mapped to 17,713 unique ICD-10-CM codes through 183,484 associations. As we did with the DO, we used this independent mapping source (henceforth, the *IHTSDO mapping project*) to discover MeSH associations with ICD-10-CM and SNOMED CT that are not included in the UMLS Metathesaurus. For this purpose, we downloaded its latest version (US1000124_20180901) and mapped the SNOMED CT and ICD-10-CM codes to MeSH descriptors using the UMLS Metathesaurus. Then, we obtained the connections of MeSH descriptors to ICD-10-CM codes via SNOMED CT codes, and the connections of MeSH descriptors to SNOMED CT codes via ICD-10-CM codes, as shown in Figure 1. Finally, in the Results section we compared these connections with those present in UMLS.

**Figure 1.**
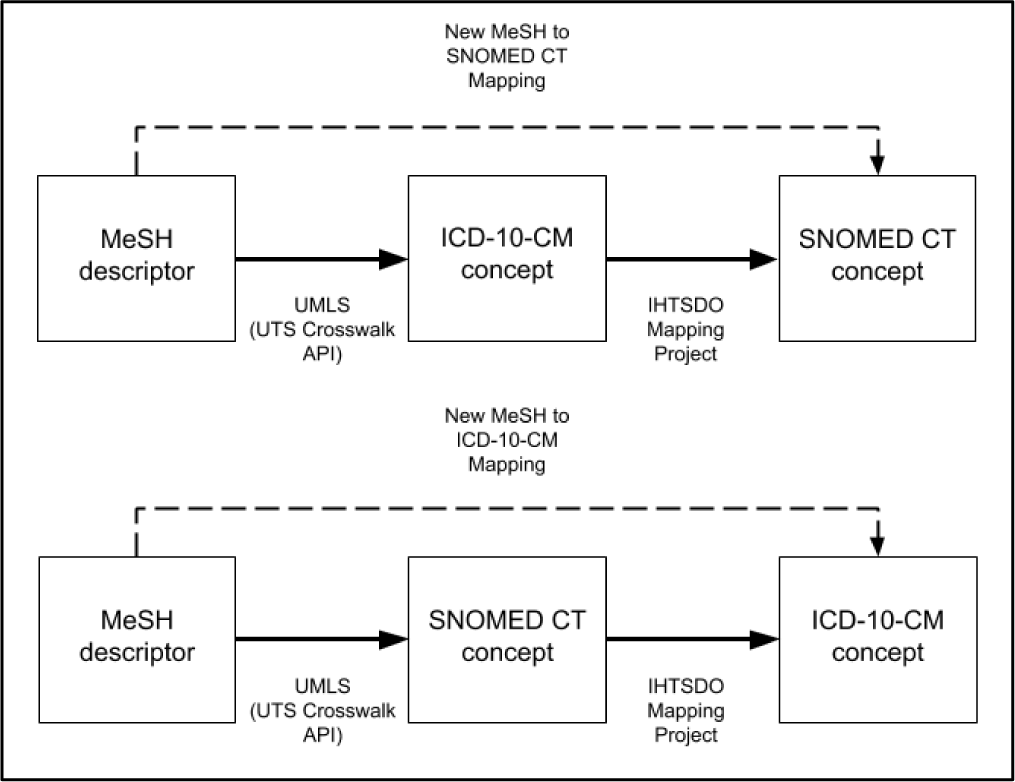
Finding new mappings of MeSH descriptors to ICD-10-CM and SNOMED CT concepts through the IHTSDO mapping project.

## III. RESULTS

### A. Missing UMLS mappings found in the DO

We found 227 mappings between 106 unique MeSH descriptors and ICD-10-CM concepts in the DO that are not present in the UMLS Metathesaurus. Of these connections, 81 correspond to MeSH descriptors that don’t contain any ICD-10-CM mapping in UMLS. For example, DOID:0002116 (*Pterygium*) relates MeSH descriptor D011625 with ICD-10-CM terms H11.00 (*Unspecified pterygium of eye*) and H11.009 (*Unspecified pterygium of unspecified eye*). These mappings are not included in the 2018AB release of UMLS.

Regarding SNOMED CT, we found 258 mappings in the DO connecting this vocabulary with 74 unique MeSH descriptors, that are not contained in the latest version of UMLS. Of these connections, 33 correspond to MeSH descriptors that don’t contain any SNOMED CT mapping in UMLS. For instance, DOID:0050477 (*Liddle Syndrome*) relates MeSH descriptor D056929 with SNOMED CT codes 190901004, 237493003, 707749005 and 71275003. These mappings are not included in the 2018AB release of UMLS. Supplementary files contain the complete list of DO resources with the MeSH to ICD-10-CM and MeSH to SNOMED CT mappings, respectively, missing in UMLS.

### B. Missing UMLS mappings found in the IHTSDO mapping project

We found 6,817 mappings between 2,549 unique MeSH descriptors and ICD-10-CM codes via SNOMED CT that are not included in the UMLS Metathesaurus. Of these connections, 1,296 correspond to MeSH descriptors that don’t contain any ICD-10-CM mapping in UMLS. For example, MeSH descriptor D037061 (*Metatarsalgia*) is connected via SNOMED CT 10085004 to ICD-10-CM codes M77.42 (*Metatarsalgia, left foot*) and M77.41 (*Metatarsalgia, right foot*). In UMLS version 2018AB, this MeSH descriptor is related only to ICD-10-CM codes M77.4 (*Metatarsalgia*) and M77.40 (*Metatarsalgia, unspecified foot*).

As for SNOMED CT, we found 34,892 mappings connecting this vocabulary with 1,924 unique MeSH descriptors via ICD-10-CM codes, that are not present in the latest version of UMLS. Of these connections, 28 correspond to MeSH descriptors that don’t contain any SNOMEDCT mapping in UMLS. For instance, MeSH descriptor D000067836 (*Hoarding disorde*r) is mapped via ICD-10-CM code F42.3 to SNOMED CT 248025009 (*Hoarding*). This MeSH descriptor is not related to any SNOMED CT code in the latest version of the UMLS Metathesaurus.

Supplementary files contain the MeSH to ICD-10-CM and MeSH to SNOMED CT mappings, respectively, extracted from the SNOMED CT to ICD-10-CM mapping project that are not included in UMLS.

## IV. DISCUSSION

The results described in the previous section show that it is possible to complete at least a part of the missing UMLS mappings of MeSH disease terms to ICD-10-CM and SNOMED CT codes by combining alternative expert-curated sources. In the case of ICD-10-CM, of the 2,459 MeSH descriptors not associated with this vocabulary in UMLS, a total of 1,322 (53.76%) were found. The other new mappings correspond to 1,266 MeSH descriptors that are already associated with at least one ICD-10-CM code in UMLS. Figure 2 illustrates the distribution of the mappings found through each source with respect to the mappings not available in UMLS.

**Figure 2.**
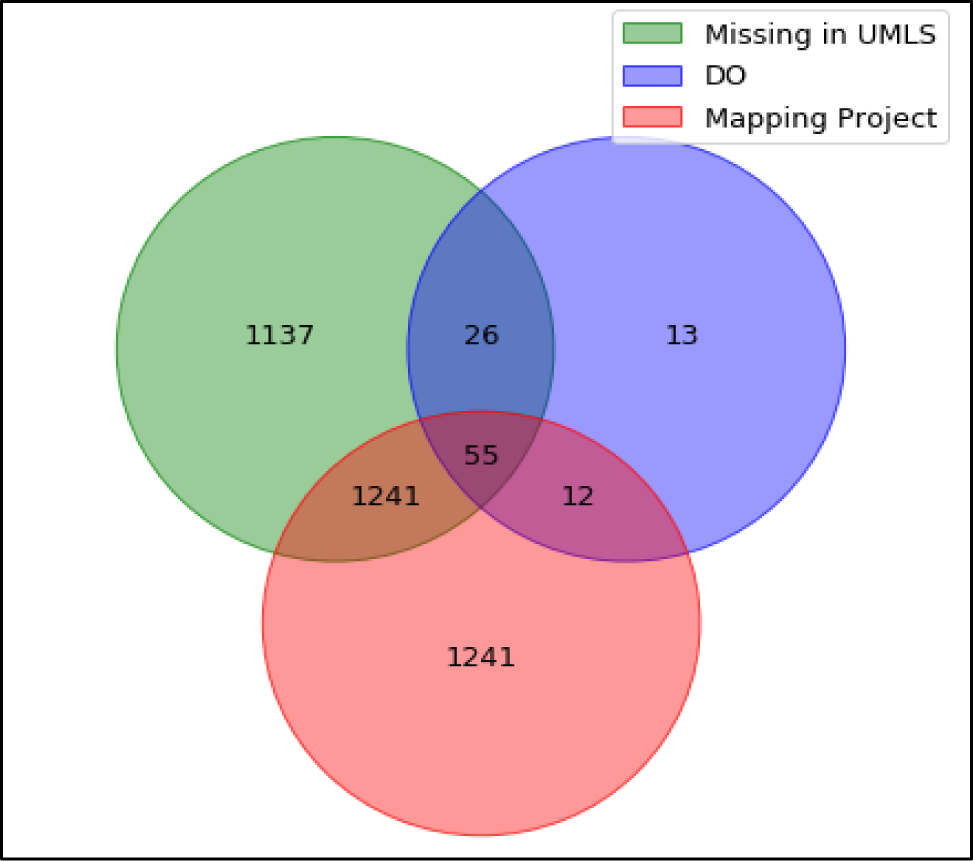
Distribution of MeSH descriptor mappings to ICD-10-CM codes found through the DO and the Mapping Project respect to those missing in the UMLS.

Regarding SNOMED CT, of the 704 MeSH descriptors not associated with this vocabulary in UMLS, a total of 56 (7,95%) were found. The other new mappings found correspond to 1,922 MeSH descriptors that are already associated with at least one SNOMED CT code in UMLS. Figure 3 illustrates the distribution of the mappings found through each source with respect to the mappings not available in UMLS.

**Figure 3.**
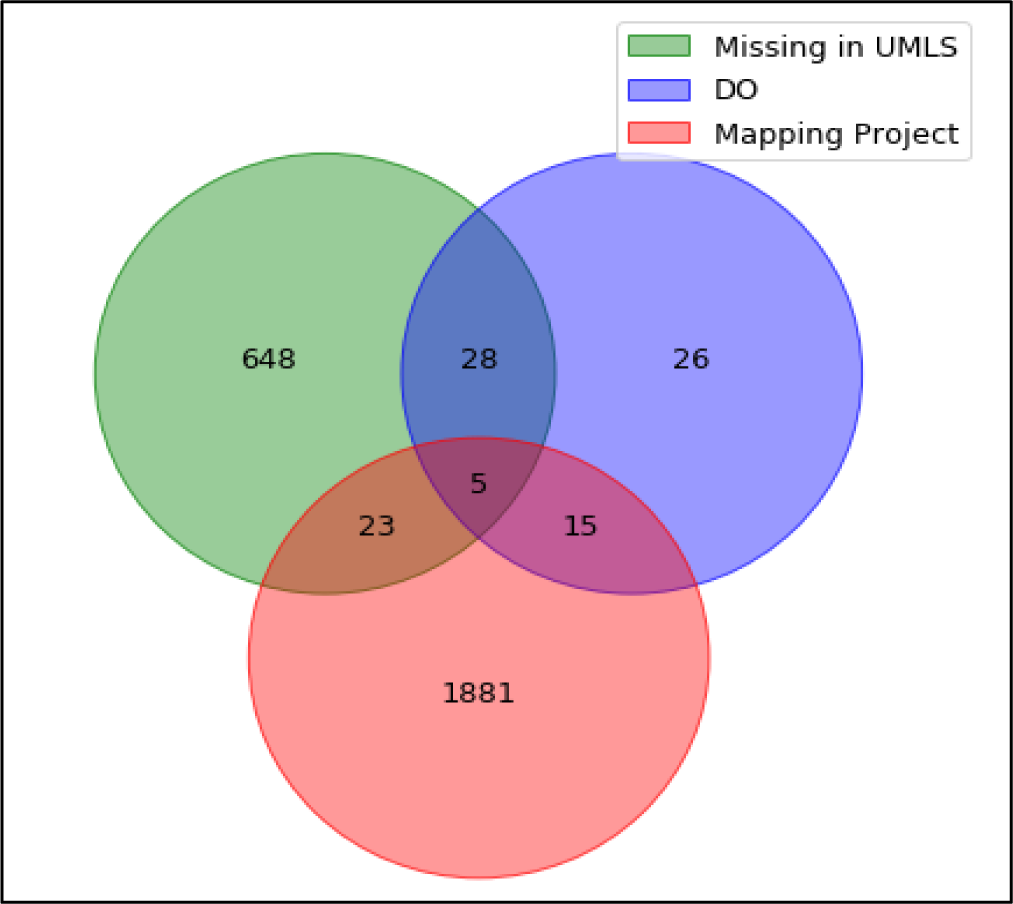
Distribution of MeSH descriptor mappings to SNOMED CT codes found through the DO and the Mapping Project respect to those missing in the UMLS.

Both in the case of ICD-10-CM and SNOMED CT, the overlap of new mappings found simultaneously in the DO and the IHTSDO mapping project is small, which indicates a low redundancy in their data.

## V. CONCLUSIONS

Improving our disease understanding requires aggregating more and more information sources. It is therefore imperative to have resources that interconnect these sources, mapping their different codes. UMLS is playing a very important role in this area, although there is still work to be done. Despite the various initiatives developed to automate the process, completing the mapping between vocabularies is costly and expert validation is essential.

In this article we have proposed the exploitation of other existing reviewed resources, such as the DO and the mapping project between SNOMED CT and ICD-10-CM by the IHTSDO, to complete the mappings between MeSH disease descriptors and disease codes of SNOMED CT and ICD-10-CM that are not yet available in UMLS. Combining these resources with the UMLS tools, we have efficiently obtained new mappings for 4,653 MeSH descriptors to these vocabularies. Of the missing connections of MeSH with ICD-10-CM and SNOMED CT, near 54% and 8% of them were resolved, respectively. All the results have been shared for verification, reuse and expansion in future projects.

While the study has focused on the mapping of disease descriptors in MeSH with SNOMED CT and ICD-10-CM, the same approach could be used for other classification systems such as OMIM or NCI. It is also proposed as future work to study the mapping coverage of the Supplementary Concept Records (SCRs) of MeSH in UMLS. The SCRs are used to index terms in the literature that are not currently MeSH headings. There are currently over 230,000 SCR records with over 505,000 SCR terms. In the case of diseases, SCRs are frequently used with rare diseases, which makes their study especially interesting in terms of mapping with classification systems such as Orphanet.

## ACKNOWLEDGMENT

This paper is supported by European Union’s Horizon 2020 research and innovation programme under grant agreement No. 727658, project IASIS (Integration and analysis of heterogeneous big data for precision medicine and suggested treatments for different types of patients).

## Footnotes

1 https://github.com/pantapps/cbms2019

